# Insulated expression of periplasmic uricase in *E. coli* Nissle 1917 for the treatment of hyperuricemia

**DOI:** 10.1101/2022.04.17.488604

**Authors:** Lina He, Wei Tang, Ling Huang, Wei Zhou, Shaojia Huang, Linxuan Zou, Lisha Yuan, Dong Men, Shiyun Chen, Yangbo Hu

**Author notes:** For correspondence: D.M., S.C. or Y.H. These authors contributed equally to this work (Lina He, Wei Tang).

## Abstract

Hyperuricemia is a prevalent disease worldwide that is characterized by elevated urate levels in the blood owing to purine metabolic disorders, which can result in gout and comorbidities. As approximately one-third of urate is excreted by the small intestine and cleared by intestinal microorganisms, modulating the gut microbiota could be an attractive approach for hyperuricemia and gout treatment. In this study, we engineered a probiotic *E. coli* Nissle 1917 (EcN) strain, EcN C6, which expresses periplasmic uricase at an “insulated site”, for urate degeneration. Oral administration of EcN C6 successfully alleviated hyperuricemia, related symptom and gut microbiota in a purine-rich food-induced hyperuricemia rat model and a *uox*-knockout mouse model. Importantly, the expression of periplasmic uricase in the insulated site did not influence the probiotic properties or global gene transcription of EcN, suggesting that EcN C6 is a safe, effective and low cost therapeutic candidate for hyperuricemia treatment.

## Introduction

Hyperuricemia is a highly prevalent disease characterized by elevated urate (uric acid) levels in the blood owing to purine metabolic disorders. This disease is also the most important risk factor for the development of gout ^1,2^. The prevalence of hyperuricemia ranges from 10% to 20% in developed countries ^3-5^. Consequently, the incidence of gout is 5% in the US, 4.75% in Europe, and 3.8% in Australia ^4-7^. Importantly, many hyperuricemic patients are asymptomatic and thus clinically neglected; however, diseases caused by hyperuricemia cannot be ignored ^5^. Increasing evidence suggests that hyperuricemia is a risk factor for the development of a variety of comorbidities, including hypertension, diabetic renal disease, obesity, metabolic syndrome, fatty liver, and cardiovascular disease ^2,5,8-10^. Therefore, hyperuricemia remains to be a global public health issue.

Urate is a key product of the purine metabolic pathway and is highly conserved in living organisms ^11^. In most species, urate is metabolized to a more soluble compound called allantoin by urate oxidase (uricase), and is further degraded to urea or ammonia ^12^. In contrast, the uricase gene found in ancestral apes has been silenced in humans owing to evolutionary events; thus urate is the final product of the purine metabolic pathway ^12,13^. Approximately 2/3 of urate in the human body is excreted by renal urate transporters (such as GLUT9 and URAT1), while the remaining 1/3 is transported by the ABCG2 transporter in the small intestine (via the extra-renal excretion pathway) and cleared by intestinal microorganisms via a process known as uricolysis ^14-16^. The overproduction or underexcretion of urate is the main cause of hyperuricemia. Therefore, traditional pharmacological urate-lowering therapies (ULTs) target urate generation (xanthine oxidase inhibitors) ^6,17^ or renal urate excretion (uricosurics) ^18^ or directly increase urate degradation (uricase) ^19,20^. However, these drugs have potential severe adverse effects and are not recommended for a large proportion of patients with asymptomatic hyperuricemia ^21^.

The intestinal tract plays an increasing role in urate excretion, particularly in patients with chronic kidney disease (CKD) whose renal elimination of urate is impaired ^19,22^. The reduction of extrarenal urate excretion is a common cause of hyperuricemia in patients with CKD ^23,24^. According to previous studies, dysbiosis of intestinal flora exists in patients with gout and serum urate (sUA) levels are associated with gut microbiome changes ^25,26^. Therefore, modulation of the gut microbiota is an alternative approach to hyperuricemia treatment ^27^. Appropriate supplementation of probiotics plays a role in urate-lowering by regulating the intestinal flora ^3,28^. Moreover, urate concentration in intestinal has been found to be positively related to sUA, and oral administration of uricase can reduce sUA in hyperuricemic rats and urate oxidase-deficient mice ^19,21,29^, suggesting that the use of engineered probiotics expressing functional uricase is an attractive strategy for the treatment of hyperuricemia.

### Escherichia coli

Nissle 1917 (EcN) is a probiotic with superior intestinal adaptation ^30,31^ that has been modified as a bacterial “living factory” for various applications. As a result, several recombinant strains have been used in clinical trials ^32,33^. Although many of the current modifications are constructed by expressing exogenous genes to endow new functions ^32,34-36^, it is unclear whether these edits will affect their probiotic characteristics, as global gene expression might be affected in these engineered strains ^32,37^. A strategy to express exogenous genes with desired functions without affecting background gene expression would strengthen the clinical applications of recombinant EcN strains ^38^.

In this study, we engineered a strain called EcN C6 with insulated expression of uricase from *Cyberlindnera jadinii*. We characterized that the fusion of the uricase gene with the TAT signal peptide FtsP is essential for efficient degradation of urate by the strain. Further, we demonstrated the effects of EcN C6 in alleviating hyperuricemia and related symptom and restoring the gut microbiota disturbed by hyperuricemia in rat and mouse models. Collectively, our data suggest that EcN C6 is a safe and effective therapeutic candidate for hyperuricemia.

## Results

### EcN expressing uricase in the periplasmic space effectively degrades urate *in vitro*

To engineer a strain with urate degradation activity, we designed a strain based on EcN to express functional uricase, which catalyzes the oxidation of urate to allantoin (Fig. 1a). To our surprise, cytoplasmic expression of uricase in EcN using a P15A originated plasmid showed markedly low urate degradation activity *in vitro* (Supplementary Fig. 1a). To explore whether the expression location of uricase is important, we fused the uricase gene with different secretion signal peptides, but most of these signal peptides could not enhance uricase activity (Supplementary Fig. 1a). Interestingly, when the uricase gene was fused with the TAT secretion signal peptide, FtsP (SufI), which drives uricase into the bacterial periplasmic space (Fig. 1b) ^39^, urate degradation efficiency was remarkably improved (Supplementary Fig. 1b). Consistent with the location of the periplasmic space, the supernatant from FtsP-uricase strain did not show uricase activity, suggesting that there was no uricase leakage into the supernatant (Supplementary Fig. 1c).

**Fig. 1.**
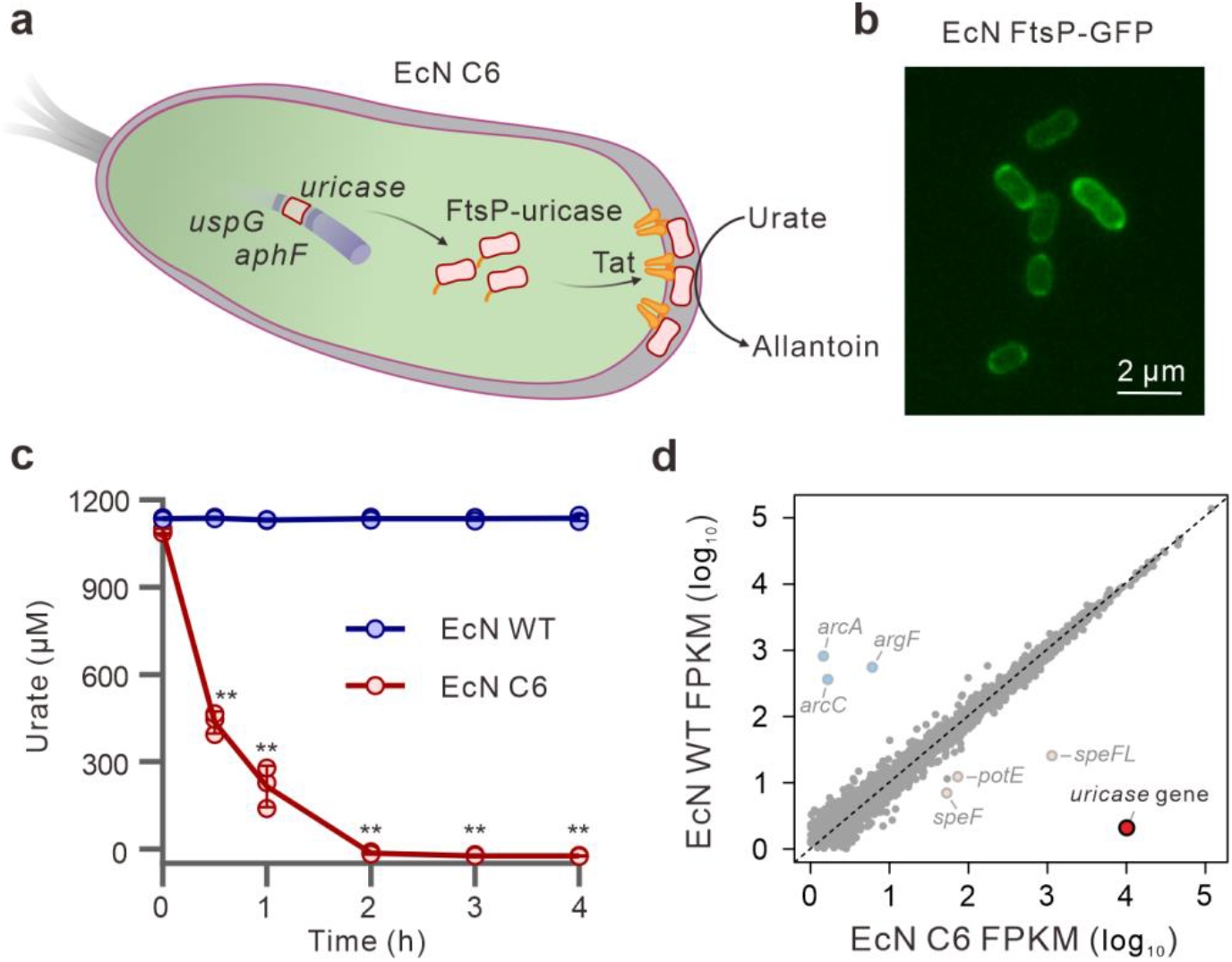
Periplasmic expression of uricase in engineered EcN C6 to degrade urate *in vitro*. **a** Schematic of the engineered strain, EcN C6, with periplasmic expression of uricase by degradation of urate to allantoin. **b** Periplasmic localization of GFP fused with the FtsP signal peptide. **c** Urate degradation activity of EcN C6 *in vitro*. **d** Linear plots of gene expression in the EcN WT and C6 strains as tested by RNA-seq analysis. Data represent measurements from three independent bacterial cultures; bars indicate the mean ± SD based on two-tailed unpaired Student’s t-test. (**P < 0.01).

To minimize the influence of uricase expression on global gene expression as well as probiotic properties of EcN, we selected a site located between *uspG* and *ahpF* (Supplementary Fig. 2a) for exogenous gene integration in the EcN genome, as this non-coding region showed markedly low gene transcription comparing with the surrounding coding regions based on published RNA-sequencing data ^40-42^. The 3’-end of both *uspG* and *ahpF* showed two opposite ρ-independent terminator structures, T1 and T2, respectively (Supplementary Fig. 2b), suggesting that the region between these two terminators would be an ideal “insulated site”. At this site, we inserted a uricase-expressing cassette containing a synthesized σ^70^-dependent promoter, a coding region carrying the FtsP signal peptide in fusion with the uricase gene, and an *rrnB* terminator (*rrnB*T), into this insulated site to obtain an engineered strain named EcN C6 (Fig. 1a, Supplementary Fig. 2a). Compared with EcN wild-type (WT), EcN C6 degraded urate *in vitro* within two hours (Fig. 1c), but showed similar phenotypes as demonstrated by its growth curve (Supplementary Fig. 3a) and also the ability to kill *Salmonella* Typhimurium LT2 under low iron conditions (Supplementary Fig. 3b), suggesting that the probiotic characteristics of EcN C6 were not affected by functional uricase expression. Transcriptomic analysis showed only six genes related to the arginine biosynthesis pathway were significantly affected by uricase expression as shown by global transcriptional profiling (Fig. 1d). Together, we successfully engineered a recombinant EcN strain with insulated expression of periplasmic uricase to degrade urate *in vitro*.

### EcN C6 ameliorates hyperuricemia disease in rat model

We applied a purine-rich food-induced hyperuricemia rat model to determine whether EcN C6 can degrade urate *in vivo*. Briefly, we first treated specific pathogen-free (SPF) Sprague-Dawley (SD) rats with purine-rich food for 21 days to induce hyperuricemia. Subsequently, the rats were orally administered EcN WT or C6 with purine-rich food (Fig. 2a). In this model, the sUA levels in rats increased significantly after 21 days of purine-rich food induction (Fig. 2b). Importantly, daily treatment with EcN C6 successfully decreased sUA levels within three days after the first administration, whereas the EcN WT or gavage buffer (GB) did not decrease sUA levels in this hyperuricemic rat model (Fig. 2b).

**Fig. 2.**
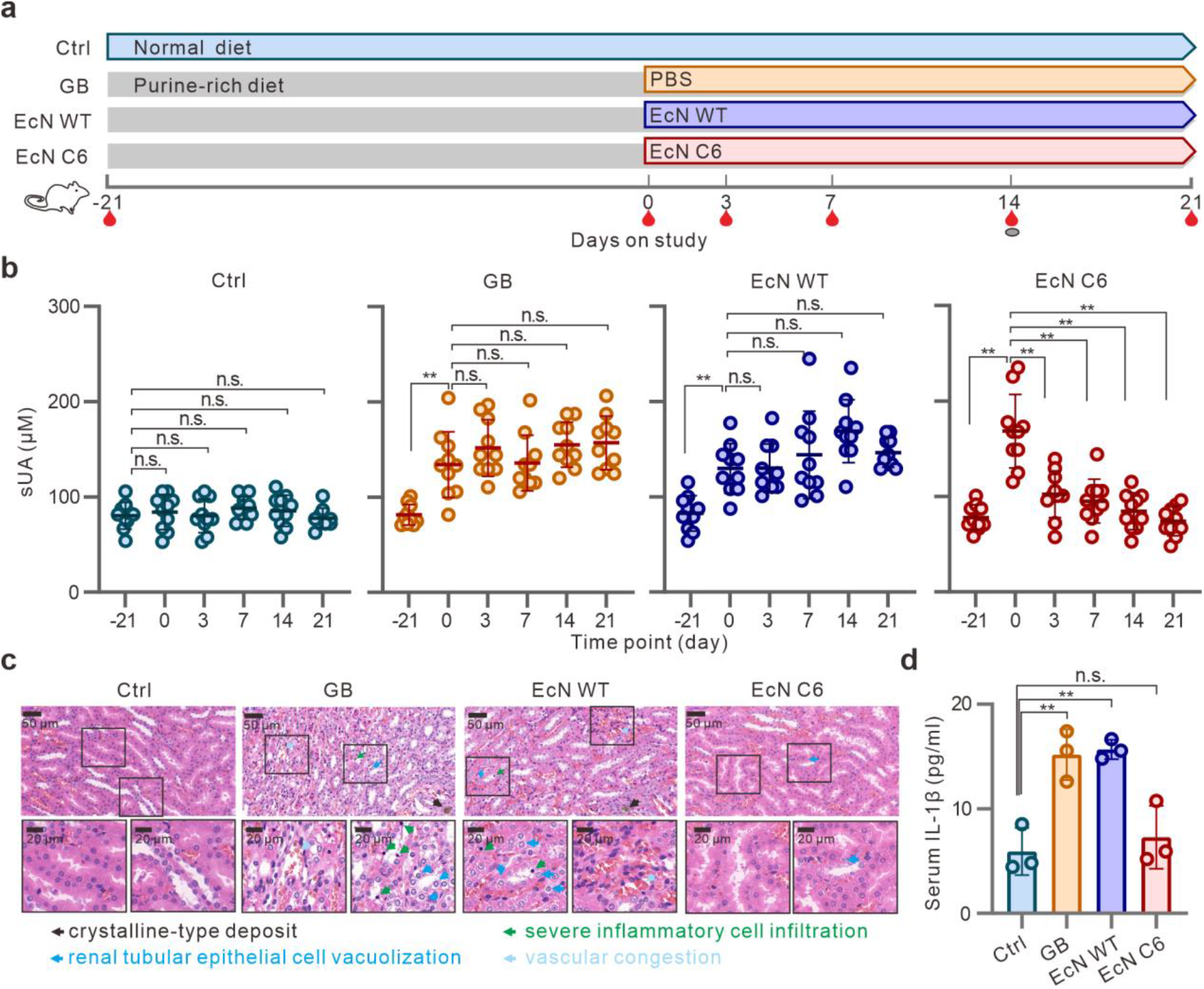
Treatment with EcN C6 decreases sUA levels and alleviates kidney damage in a rat model. **a** Schematic representation of the treatment of recombinant EcN C6 expressing uricase in a rat model of hyperuricemia. Male SD rats (n = 40) were divided into four groups (10 for each group); three quarters were pretreated with purine-rich food for 21 days to induce hyperuricemia and the remaining one quarter was used as the control with no treatment (Ctrl). The EcN WT, EcN C6 (3×10^10^ CFU for each strain), or gavage buffer (GB) was administered orally to the hyperuricemic rats for another 21 days. Purine-rich food was also provided during this treatment to maintain high sUA level. At indicated time points, bloods and feces were collected. Rats were euthanized after 21 days of treatment for kidney imaging. **b** sUA levels of purine-rich food hyperuricemia rats after treatment with different strains. **C** Representative renal tissue sections with hematoxylin and eosin staining. Scale bars, 50 μm or 20 μm. **d** IL-1β levels in rats with different treatments. Statistical analysis was performed using two-tailed unpaired Student’s t-test (**P < 0.01).

To further explore whether EcN C6 could alleviate the pathological symptoms caused by hyperuricemia, rats were euthanized 21 days after treatment. Compared with the chow diet group, the EcN C6 group displayed attenuated urate-induced inflammation, as demonstrated by renal pathology HE staining and serum IL-1β levels (Fig. 2, c and d). Groups administered GB or the EcN WT strain showed significant renal crystals, severe inflammatory cell infiltration, and large amount of vacuolation in renal tubular epithelial cells and interstitial congestion of renal tubules (Fig. 2c). Together, these data illustrate that EcN C6 possesses urate-lowering effects and alleviates hyperuricemia symptoms in a rat model, suggesting that the EcN C6 strain is applicable for the treatment of hyperuricemia.

### EcN C6 alleviates dysbiosis of the gut microbiota in hyperuricemia rats

To explore the effect of EcN C6 on gut microbes in a hyperuricemia rat model, we extracted fecal bacterial DNA from rats before and after 21 days of purine-rich food induction, as well as 14 days after EcN C6 treatment in the purine-rich food-induced hyperuricemia model. 16S rRNA gene amplicon sequencing was then employed to detect bacterial species in rats at different stages, which revealed that *Bacteroidetes* and *Firmicutes* were the two most abundant gut microbial phyla (Fig. 3a). Principal component analysis (PCA) demonstrated that the gut microbial composition changed significantly in the hyperuricemia rat model (day 0 compared with day -21) (Fig. 3b). Further, statistical analysis revealed that hyperurate induced gut flora disorders, including alterations in the contents of *Bacteroidetes, Firmicutes*, and *Proteobacteria* (Fig. 3, c-e), as well as *Verrucomicrobia* and *Deferribacteres* (Supplementary Fig. 4), whereas treatment with EcN C6 alleviated dysbiosis in the gut microbiota (Fig. 3, Supplementary Fig. 4). Overall, these results suggest that EcN C6 may balance the gut microbiota in rats with hyperuricemia.

**Fig. 3.**
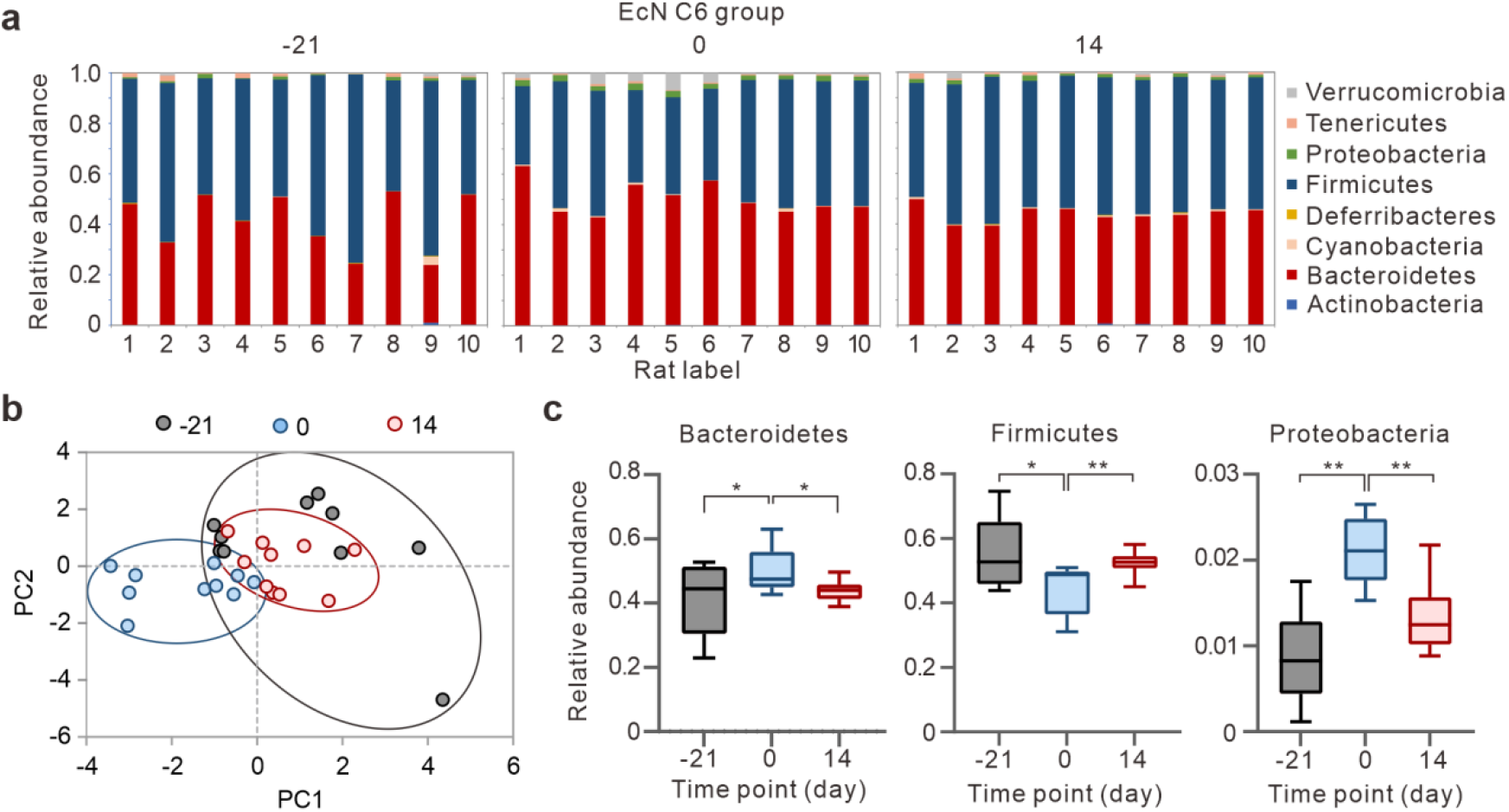
EcN C6 alleviates gut microbiota dysbiosis in hyperuricemia rats. **a** Comparison of phylum relative abundance of rats before (−21) and after (0) high-purine food induction, as well as treatment with the EcN C6 strain for 14 days (14). **b** Principal coordinates plot of the gut microbiota in rats treated with EcN C6 at different time points as in A. **c** Relative abundance of *Bacteroidetes, Firmicutes*, and *proteobacteria* in the gut microbiota of the EcN C6 group at -21, 0, and 14 days (n = 10). Statistical analysis was performed using two-tailed unpaired Student’s t-test (**P < 0.01).

### Administration of EcN C6 twice per week lowers sUA levels

To explore the effective dosage of EcN C6 for lowering the sUA level, we treated the hyperuricemic rats with different doses of EcN C6 (Fig. 4a). As expected, sUA levels in hyperuricemic rats were significantly decreased 3 days after a single-dose administration of EcN C6 but were restored after 7 days (Fig. 4b). Similarly, once per week treatment resulted in the same trend (Fig. 4b), suggesting that the once per week dose was insufficient to effectively lower the sUA levels. However, administration twice per week successfully lowered the sUA levels, which then remained at a normal level (Fig. 4b). This discrepancy was also reflected in the renal pathology HE staining and serum creatinine, urea nitrogen, and IL-1β levels of the different dosage groups (Supplementary Fig. 5). Continuous monitoring of EcN C6 colonization in feces by qPCR revealed that it could be detected within 2-3 days after administration (Fig. 4c), which is consistent with other reports ^32,35^. The twice per week dose group displayed two detecting peaks within one week (Fig. 4c), which further supports our conclusion that the maintenance of EcN C6 in the gut is important for lowering urate levels.

**Fig. 4.**
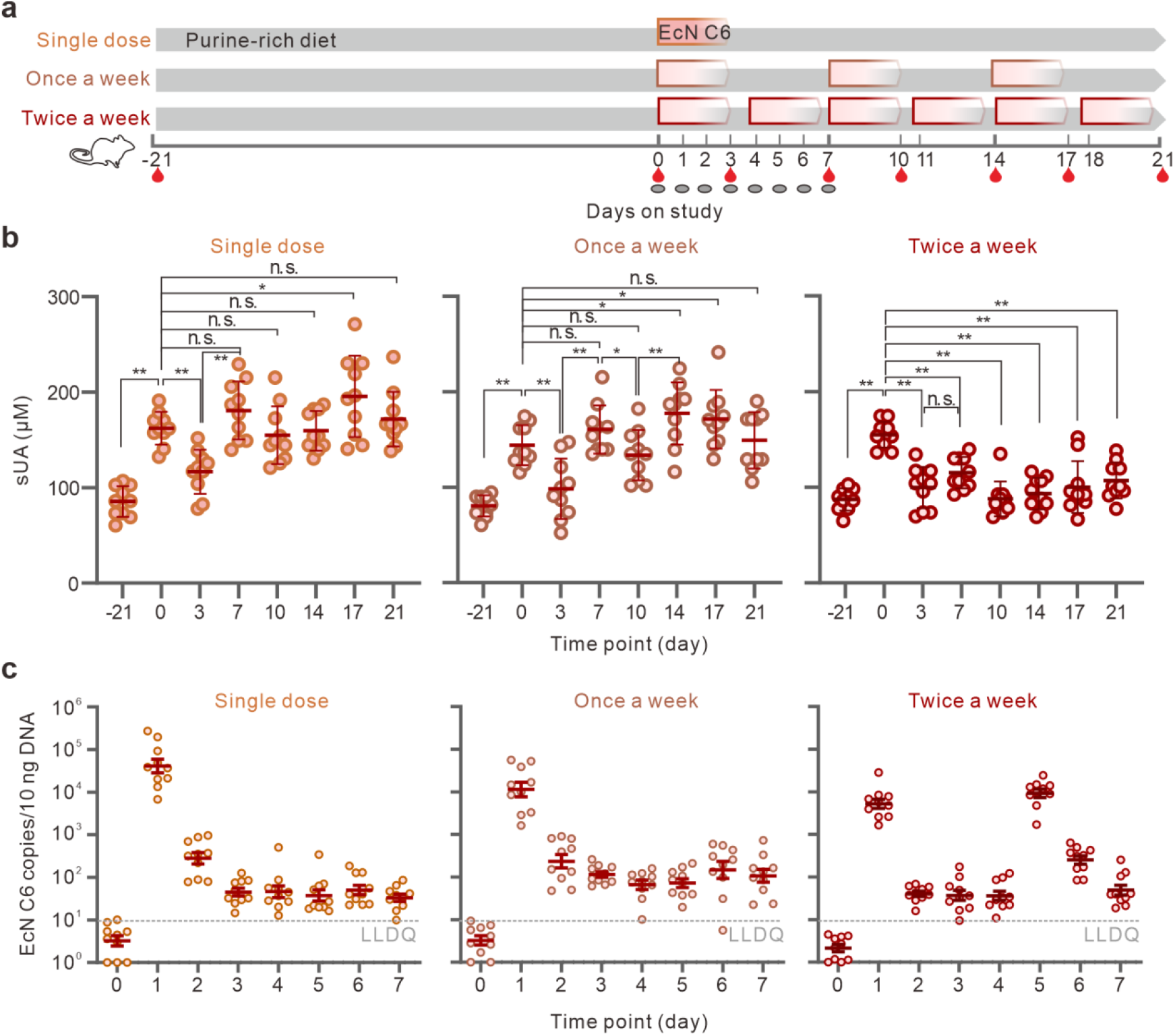
Administration of EcN C6 twice per week is required to lower sUA levels. **a** Schematic representation of the treatment of hyperuricemia rats with different doses of EcN C6. Male SD rats (n = 30) were pretreated with purine-rich food to induce hyperuricemia, and divided into three groups: Single dose group (1.2×10^11^ CFU), once per week group (3×10^10^ CFU) and twice per week group (3×10^10^ CFU). Blood samples and feces were collected at indicated time points. **b** sUA levels of hyperuricemia rats administered purine-rich food after treatment with different doses of the EcN C6 strain. **c** Detection of EcN C6 copies in feces at different time points (n = 10 for each group). Statistical analysis was performed using two-tailed unpaired Student’s t-test (**P < 0.01).

### EcN C6 alleviates hyperuricemia symptoms in a *uox*-knockout mouse model

We employed a *uox*-knockout mouse model to further confirm the effect of EcN C6 on lowering sUA levels. As knockout of the *uox* gene is detrimental ^43^, only 12 knockout mice were obtained. Stably elevated sUA, serum creatinine and urea nitrogen levels in these mice indicated that the model had been successfully established (Fig. 5a). Mice were then divided into two groups and treated with EcN WT and EcN C6 (Fig. 5b). In contrast to the WT group, the EcN C6 group had significant alleviation of the hyperuricemia indicators, including sUA, creatinine and urea nitrogen (Fig. 5c). Similar to our observations in the rat model, EcN C6 treatment lowered the inflammatory response as reflected by the renal pathology HE section (Fig. 5d) and kidney IL-1β levels (Fig. 5e). Taken together, our data suggest that EcN C6 can alleviate the symptoms of hyperuricemia in a *uox*-knockout mouse model.

**Fig. 5.**
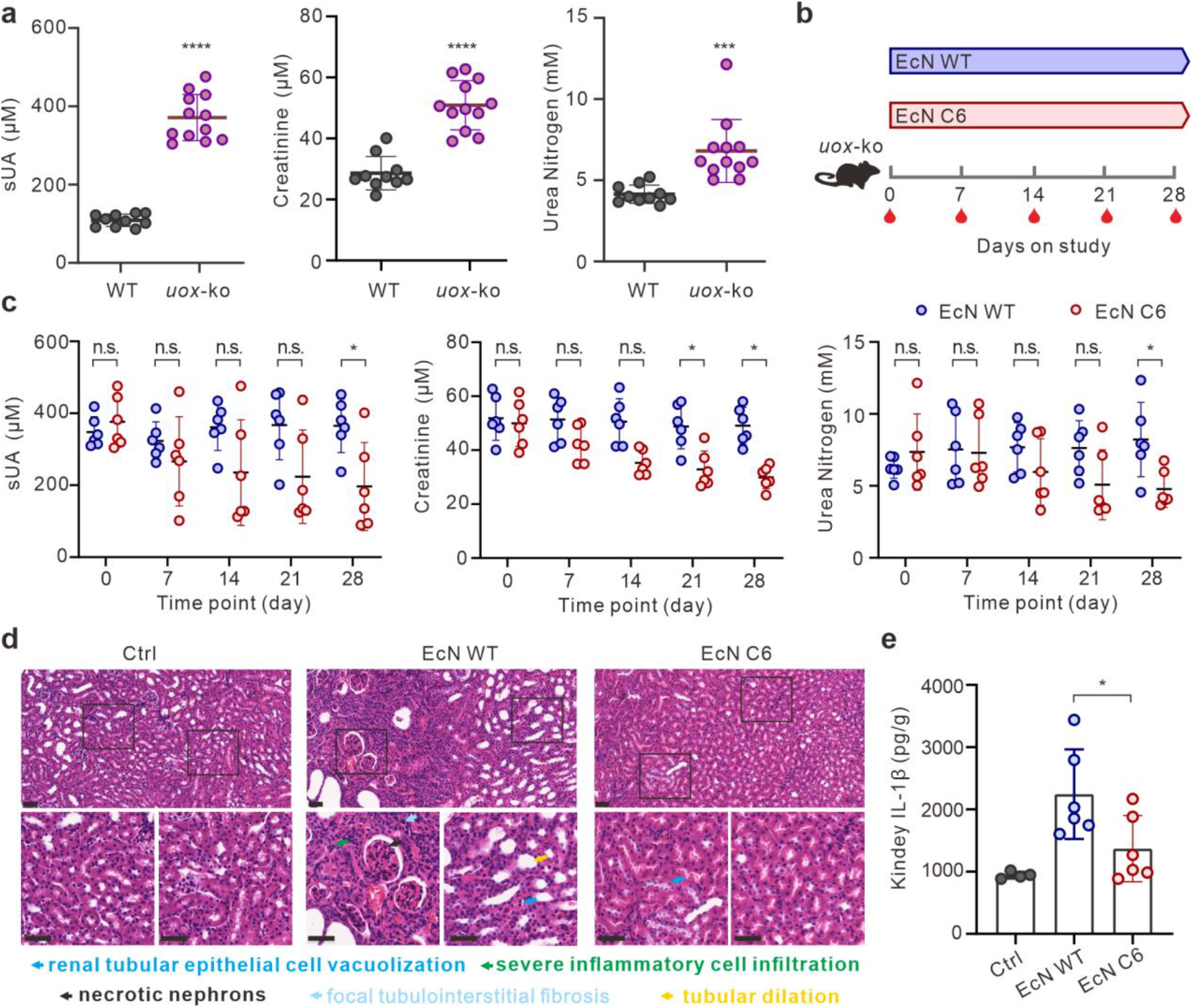
Effects of EcN C6 treatment on hyperuricemia symptoms in a *uox*-ko mouse model. **a** Serum urate, creatinine, and urea nitrogen levels of wild-type (n = 10) and *uox*-ko (n = 12) mice. **b** Schematic representation of the treatment with EcN WT (1×10^10^ CFU) or EcN C6 (1×10^10^ CFU) for one month in the *uox*-ko mouse model. Blood samples were collected at indicated time points. After treatment for 28 days, the kidneys were dissected for tissue HE staining and inflammatory factor detection. **c** Serum urate, creatinine, and urea nitrogen levels in mice administered EcN WT or EcN C6 at indicated time points (6 for each group). **d** Representative renal tissue sections with HE staining. Scale bars, 50 μm. **e** Kidney IL-1β levels in wide-type mice (Ctrl, n = 4) or *uox*-ko mice administered EcN WT or EcN C6 treatment (6 for each group). Statistical analysis was performed using two-tailed unpaired Student’s t-test (*P < 0.1; ***P < 0.001).

## Discussion

Gout is a common and challenging health issue worldwide. Despite the availability of treatments for lowering urate levels, these drugs mainly aim to inhibit urate synthesis or promote urate excretion, thereby placing a remarkable burden on the kidneys ^23,44^ and may have side effects on gut bacteria ^45-47^. Here, as summarized in Fig. 6, we successfully engineered a probiotic strain, EcN C6, with insulated expression of periplasmic uricase to directly degrade urate and alleviate the symptoms and dysbiosis of the gut microbiota caused by hyperuricemia, ultimately providing an efficient and friendly method for the rapid treatment of hyperuricemia.

**Fig. 6.**
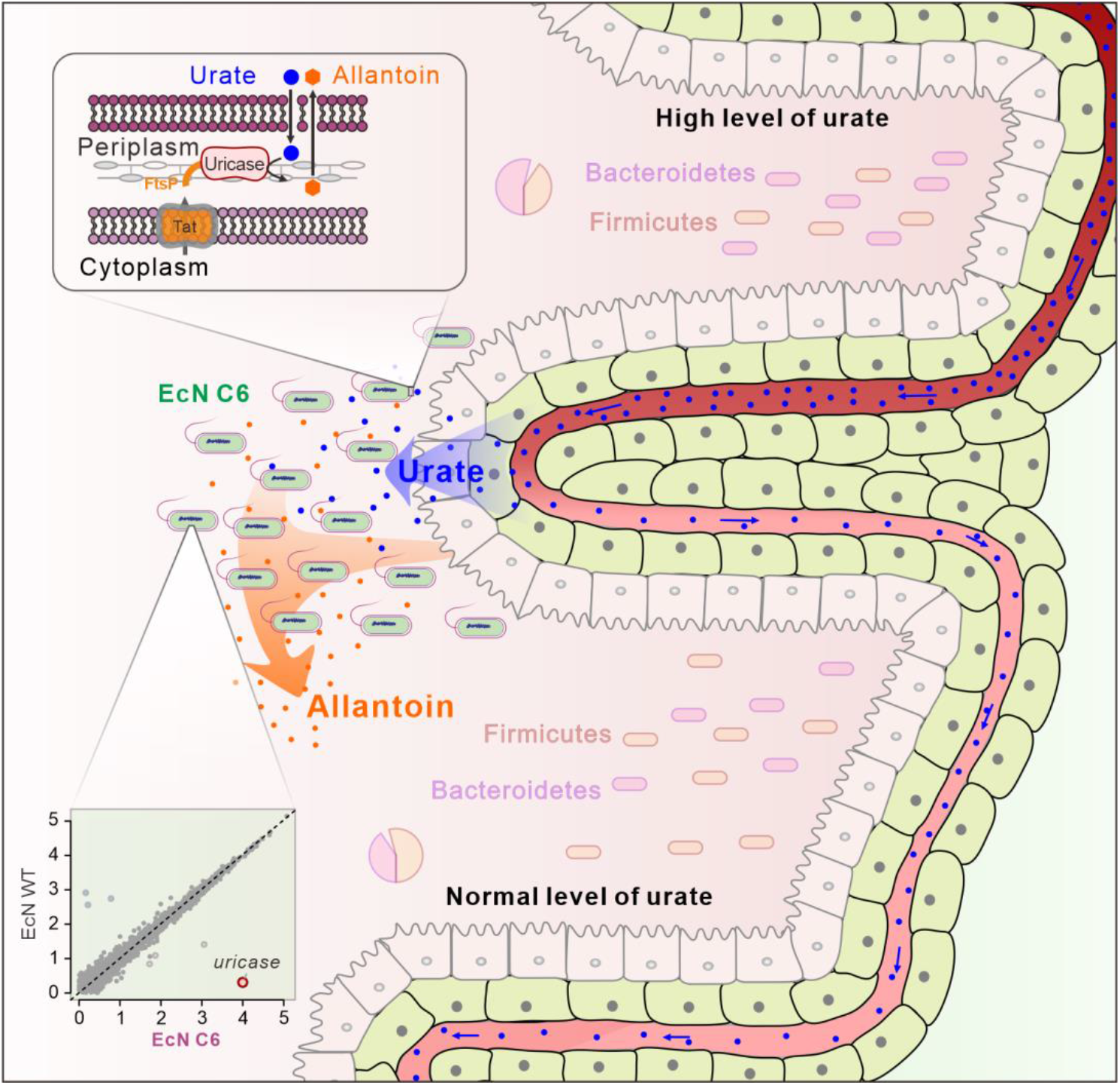
Proposed model for the action of EcN C6 strain in lowering sUA levels *in vivo*. In a hyperuricemia model, urate in the blood (blue dots) is transferred to the intestine by urate transporters. Engineered EcN C6 (green ellipses) with insulated expression (down insert) of periplasmic uricase (up insert) degrades urate into allantoin in the gut. Meanwhile, the administration of EcN C6 alleviates dysbiosis of the gut microbiota in the hyperuricemia model.

In addition to traditional urate-lowering chemical drugs, enzymes related to the degradation of urate are gaining attention, and may serve as a more direct method for hyperuricemia treatment ^48,49^. Clinical data have shown that modified uricases, such as pegolase, lablipase, and pregabalin, display excellent performance in the treatment of intractable gout disease by intravenous injection; however, their duration of action *in vivo* is limited. Notably, large amounts of supplementation can induce antibody production and are cost-effective ^50,51^. Uricases are strongly not recommended as first-line therapy by the American College of Rheumatology guidelines for the management of gout owing to their limited duration of action ^52^. To overcome the limitation, we engineered a probiotic EcN C6 expressing periplasmic uricase (Fig. 6). Probiotic EcN is the preferred microbial synthetic biology vector and can successfully colonize in the small intestine ^35^, which plays an essential role in regulating urate levels ^29^. Moreover, colonization of EcN near the epithelial cells in the small intestine allows them to easily access to urate that is transported from the blood ^29^ and obtain oxygen ^35,53^ as a necessary substrate for uricase function. This strategy has markedly extended the application of uricase for hyperuricemia treatment.

The location of the uricase expressed in EcN is essential for its ability to degrade urate. Although *aegA* and *ygfT* can degrade urate under microaerobic or anaerobic conditions in *E. coli* ^54^, in contrast to uricase, this effect was almost negligible both *in vitro* and *in vivo* (Fig. 1d and 2b). In addition, there are 10 nucleobase-ascorbate transporter (NAT) family-related proteins in *E. coli* that are responsible for transporting different forms of base metabolites. Further, the *ygfU* gene located at the inner membrane was hypothesized to import urate ^55^; however, our data showed that cytoplasmic expression of uricase in the EcN strain only slightly decreased the urate level *in vitro* (Supplementary Fig. 1b), suggesting that the efficiency of urate import may be limited by the bacterial cell membrane or the affinity of urate transporters. Therefore, we attempted to use the Sec and Tat export pathways to drive uricase out of the cytoplasm. Interestingly, in contrast to previously engineered uricase-producing *E. coli* strains ^56^, adding the Tat secretory signal peptide FtsP successfully enabled the EcN strain to exhibit sustainable degradation of urate *in vitro* and *in vivo* (Supplementary Fig. 1b, Fig. 2), which suggests that the location of uricase is essential for its activity in EcN.

Our EcN C6 design included a feature to insulate uricase expression. As a safe and common bacterial vector, EcN has been adopted for the regulation of mucosal immunity ^57,58^, metabolic diseases ^32-34^ and pathogenic infections ^59-61^. However, the expression patterns for these functional proteins are mainly based on recombinant plasmids, which fail to comply with the FDA’s requirements for live biotherapeutic products ^62^. Previous studies have attempted to insert exogenous genes near the *malEK, malPT*, and *yicS/nepI* genes in EcN ^32,63^; however, these types of genomic editing may interfere with native gene transcription ^32,38^. As the design of ultra-stable genetic editing in *E. coli* is important for living therapeutics ^38^, we characterized an “insulated site” in the EcN genome by analyzing previous transcriptome data of EcN under different conditions ^32,40,41^. This “insulated site” is flanked by two naturally appositive terminators. Our RNA-seq data revealed that this “insulated site” ensures insulated expression of the inserted uricase with the exception of six genes involved in converting NH_3_ to arginine. Therefore, our insulated expression strategy is effective and valuable for the recombinant expression of other target genes in EcN.

## Materials and Methods

### Study design

The objectives of this study were to engineer a probiotic EcN strain with insulated expression of uricase and to evaluate the therapeutic activity of this strain in murine models of hyperuricemia. We first analyzed the published transcriptome data to identify an “insulated” site located between the *uspG* and *ahpF* genes in the EcN genome. Next we applied homologous recombination to insert a cassette expressing periplasmic uricase at this insulated site to obtain an engineered strain called EcN C6. *In vitro* urate degradation assay and global RNA-seq were subsequently applied to confirm the activity and insulated expression of uricase in EcN C6, respectively. Furthermore, urate degradation activity of EcN C6 was evaluated in a purine-rich food-induced hyperuricemia rat model as well as in a *uox*-knockout mouse model. Replication and sample sizes for all experiments are described in the figure legends. Because of the exploratory nature of this study, a priori sample size estimates were not performed for the analyses described herein.

### Bacterial Strains

The bacterial strains used in this study are listed in Supplementary Table 1. *E. coli* and *S*. Typhimurium LT2 strains were grown in Luria-Bertani (LB) medium or agar (LBA) at 37 °C and supplemented with 50 μg/mL kanamycin, 100 μg/mL ampicillin, or 30 μg/mL chloramphenicol, when necessary. EcN WT and C6 were prepared in LB and the cell pellets were re-suspended in protective buffer (15% v/v glycerol, 5% w/v trehalose, and 10 mM MOPS, pH 7.3) and frozen at -80 °C until use, as previously described ^32^.

### Plasmid constructions

The plasmids and oligonucleotides used in this study are also listed in Supplementary Table 1. To express uricase fused with different signal peptides, a uricase gene from *C. jadinii* (GenBank: XM_020212619.1) with a signal peptide coding region from either *ftsP* (N-terminal 30 aa, GenBank: NP_417489.1), *ompA* (N-terminal 27 aa, GenBank: NP_415477.1), *tamA* (N-terminal 27 aa, GenBank: NP_418641.1), *lpp*-*ompA* (N-terminal 29aa of *lpp* and 46aa-159aa of *ompA*, GenBank: NP_310411.1 and NP_415477.1), *yebF* (N-terminal 118 aa, GenBank: NP_416361.2), or *inpNC* (N-terminal 211aa and C-terminal 99aa, GenBank: AF013159) was cloned into the pKT100 plasmid ^64^ using a ClonExpress II One Step Cloning Kit (Vazyme, China). The *fstP*-*gfp* expression clone was constructed in a similar manner by replacing the uricase gene with *gfp* fragment.

### Fluorescence imaging

To verify the periplasmic localization of the FtsP-GFP fusion protein, the EcN strain transformed with pKT-*ftsP*-*gfp* was grown at 37 °C in LB liquid medium containing Kan (50 μg/mL) to OD_600_∼0.3. Cells were collected by centrifugation at 5000 g for 3 min and resuspended in phosphate buffered saline (PBS). The samples were dropped onto a slide and fixed by slight heating. Fluorescence images were obtained using a fluorescence microscope (OLYMPUS).

### EcN C6 strain construction

To construct the EcN C6 strain, a fragment containing the synthesized promoter P6, a signal peptide encoded by *ftsP*, the uricase gene from *C. jadinii*, and the *rrnB*T terminator, was cloned into the pDM4 plasmid ^65^ between *uspG* and the *ahpF*. The resulting plasmid was transformed into *E. coli* S17-1 cells, which were used as the donor cells. EcN carrying the temperature-sensitive pKD46 plasmid ^66^ was used as the recipient cell. EcN C6 was constructed using conjugation ^65^ and the pKD46 plasmid in EcN C6 was removed by culturing the strain at 42 °C. The insertion of the uricase fragment into the EcN C6 strain was confirmed by PCR and DNA sequencing.

### *In vitro* urate degradation assay

Overnight cultures of the EcN strains were diluted 1:100 with 3 mL fresh LB liquid medium, incubated at 37 °C until OD _600_∼0.6 (0.3 mM IPTG was added when necessary), and continually incubated until OD_600_∼1.0. Cell pellets were collected by centrifugation and resuspended in an equal volume of MU medium as previously described ^15^. Urate concentration in the medium was monitored using A_293_ absorption with NanoDrop One (Thermo Fisher Scientific, USA) at the indicated time points ^15^. The standard curve between the A_293_ values and urate concentrations was examined to quantify the urate concentrations in the MU medium.

### Growth assay of EcN C6

Freshly streaked EcN WT and EcN C6 colonies were inoculated with 5 mL of LB and grown overnight at 37 °C. Cultures were transferred at a ratio of 1:100 into fresh 5 mL of LB and grown for 12 h at 37 °C. The OD_600_ was measured at the indicated time (BioTek). Competitive growth for EcN and *S*. Typhimurium LT2 under iron-rich or iron-limiting conditions was performed as previously described ^61^.

### RNA-seq analyses

To characterize the transcriptional profiles of EcN WT and EcN C6 strains, triplicate cultures of each strain were incubated overnight in LB broth at 37 °. Overnight cultures were then diluted 100-fold in 4 mL LB broth and incubated for 2.5 h. RNA was extracted using TRIzol reagent, as described in the manufacturer’s protocol (Invitrogen, USA). rRNA in the extracted RNA was removed using a Ribo-off rRNA Depletion Kit (Vazyme, China). The RNA library was constructed using the NEBNext® Ultra II RNA Library Prep Kit for Illumina (NEB, USA) and sequenced using the Illumina HiSeq X Ten platform.

### Hyperuricemia rat model

Two-week-old SPF SD rats were purchased from Beijing Vital River Laboratory Animal Technology Co., Ltd. After acclimatization for one week, rats were divided into groups (10 per group) and were either fed with high-purine food containing maintenance powder, 10% yeast (OXIFOD) and 0.1% adenine (Sangon, China) to induce hyperuricemia ^67^, or normal chow as a control. After induction for three weeks, rats were administered 1 mL of gavage buffer (GB group), EcN wild-type (EcN group, 3×10^10^ CFU/day), or EcN C6 (EcN C6 group, 3×10^10^ CFU/day) for three weeks. To explore the minimum dose of EcN C6, similar approaches were employed, except that 1.2×10^11^ CFU was used for the single-dose group. Serum samples were collected at the indicated time points to test sUA, creatinine, and urea nitrogen levels using commercial kits, according to the manufacturer’s instructions (Jiancheng, China). After treatment with these strains for 21 days, two rats in each group were euthanized by slow asphyxiation with CO_2_. The left kidney was dissected for tissue HE staining.

### *Uox*-knockout mouse model

Conventional SPF C57BL/6J^*uox/uox*^ (*uox*-ko) mice purchased from the Shanghai Model Organisms Center Inc. (SMOC) were maintained and bred at the Center for Animal Experiments at the Wuhan Institute of Virology. Allopurinol (90 μg/mL) was added to enhance the survival of newborns when administered to the mother and withdraw one week before the experiment. Mice were divided into two groups of equal sex and age. In this model, the EcN WT or C6 strain was administered daily by oral gavage. Serum was collected at the indicated time points after treatment for 28 days. Mice were euthanized by slow asphyxiation with CO_2_. The left kidney was dissected for HE staining, and the right kidney was used for tissue homogenization to determine the inflammatory factors.

### ELISA

Serum was isolated from the blood of rats and mice by low-speed centrifugation (1000 g, 10 min). Suspensions from ground kidney samples were collected by low-speed centrifugation (2000 g, 15 min). To detect the IL-1β in serum or kidney, the samples were analyzed using a rat or mouse IL-1β ELISA Kit (Neobioscience, China) following the manufacturer’s instructions.

### 16s rRNA library preparation and sequencing

Feces collected from hyperuricemic rats were frozen at -80 ° until use. DNA was extracted using the E.Z.N.A. Stool DNA Kit (OMEGA, USA) following the manufacturer’s instructions. The 16S rRNA gene (V4 region) was amplified by two-step PCR enrichment using barcodes for multiplexing ^68^. Pooled DNA was purified using AMpure XP beads (Beckman, USA). DNA libraries were constructed using the NEBNext Ultra II FS DNA Library Prep kit (NEB, USA) and sequenced using the Illumina HiSeq X Ten platform.

### Quantification of EcN C6 colonization

To quantify the colonization of EcN C6, qPCR was performed to determine the copy numbers of the EcN *fimA* gene in 10 ng of fecal genomic DNA using iTaq Universal SYBR Green Supermix (Bio-Rad, USA). Standard curves were constructed by quantitatively testing 10^8^, 10^7^, 10^6^, 10^5^, 10^4^, 10^3^, 10^2^, 10^1^, and 10^0^ copies of EcN C6 genomic DNA according to a previously described protocol ^3^. All measurements were performed in triplicate.

### Statistical analysis

Statistical significance between two groups was analyzed by unpaired Student’s t-test (two-tailed) using GraphPad Prism 8 or the R package (version 3.2.2).

### Animal Ethics

The animal experiments were approved by the Experimental Animal Ethics Committee of the Wuhan Institute of Virology, Chinese Academy of Sciences (Ethics No. WIVA14201901).

### Data and materials availability

RNA-sequencing and 16s rRNA gene sequencing reads were submitted to the NCBI Sequence Read Archive (SRA) under accession: PRJNA818111 and PRJNA818085, respectively.

## Supporting information

Combined supplementary file

## Acknowledgments

We thank the Center for Animal Experiment in Wuhan Institute of Virology for assisting with the rat and mouse experiments. This study was supported by the Young Top-notch Talent Cultivation Program of Hubei Province (to Y.H.).

## Author contributions

Conceptualization: L.H., D.M., Y.H.

Methodology: L.H., W.T., W.Z., Y.H.

Investigation: L.H., W.T., L.H., S.H., L.Z., L.Y.

Visualization: L.H., W.Z., Y.H.

Funding acquisition: Y.H.

Project administration: D.M., S.C., Y.H.

Supervision: S.C., Y.H.

Writing – original draft: L.H., S.C., Y.H.

Writing – review & editing: L.H., D.M., S.C., Y.H.

## References

1 Dalbeth, N., Merriman, T. R. & Stamp, L. K. Gout. Lancet 388, 2039–2052 (2016).

2 Dalbeth, N., Gosling, A. L., Gaffo, A. & Abhishek, A. Gout. Lancet 397, 1843–1855 (2021).

3 Wu, Y. et al. Limosilactobacillus fermentum JL-3 isolated from “Jiangshui” ameliorates hyperuricemia by degrading uric acid. Gut Microbes 13, 1–18 (2021).

4 Singh, G., Lingala, B. & Mithal, A. Gout and hyperuricaemia in the USA: prevalence and trends. Rheumatology (Oxford) 58, 2177–2180 (2019).

5 Joosten, L. A. B., Crisan, T. O., Bjornstad, P. & Johnson, R. J. Asymptomatic hyperuricaemia: a silent activator of the innate immune system. Nat Rev Rheumatol 16, 75–86 (2020).

6 Anderson, I. J., Davis, A. M. & Jan, R. H. Management of Gout. JAMA 326, 2519–2520 (2021).

7 Kuo, C. F., Grainge, M. J., Zhang, W. & Doherty, M. Global epidemiology of gout: prevalence, incidence and risk factors. Nat Rev Rheumatol 11, 649–662 (2015).

8 Richette, P. & Bardin, T. Gout. Lancet 375, 318–328 (2010).

9 Grassi, D. et al. Chronic hyperuricemia, uric acid deposit and cardiovascular risk. Curr Pharm Des 19, 2432–2438 (2013).

10 Yip, K., Cohen, R. E. & Pillinger, M. H. Asymptomatic hyperuricemia: is it really asymptomatic? Curr Opin Rheumatol 32, 71–79 (2020).

11 Hayashi, S., Fujiwara, S. & Noguchi, T. Evolution of urate-degrading enzymes in animal peroxisomes. Cell Biochem Biophys 32 Spring, 123–129 (2000).

12 Wu, X. W., Lee, C. C., Muzny, D. M. & Caskey, C. T. Urate oxidase: primary structure and evolutionary implications. Proc Natl Acad Sci U S A 86, 9412–9416 (1989).

13 Kratzer, J. T. et al. Evolutionary history and metabolic insights of ancient mammalian uricases. Proc Natl Acad Sci U S A 111, 3763–3768 (2014).

14 Sorensen, L. B. Role of the intestinal tract in the elimination of uric acid. Arthritis Rheum 8, 694–706 (1965).

15 Rouf, M. A. & Lomprey, R. F., Jr. Degradation of uric acid by certain aerobic bacteria. J Bacteriol 96, 617–622 (1968).

16 Matsuo, H. et al. Common defects of ABCG2, a high-capacity urate exporter, cause gout: a function-based genetic analysis in a Japanese population. Sci Transl Med 1, 5ra11 (2009).

17 White, W. B. et al. Cardiovascular safety of febuxostat or allopurinol in patients with gout. N Engl J Med 378, 1200–1210 (2018).

18 Silverman, W., Locovei, S. & Dahl, G. Probenecid, a gout remedy, inhibits pannexin 1 channels. Am J Physiol Cell Physiol 295, C761–767 (2008).

19 Pierzynowska, K. et al. Oral treatment with an engineered uricase, ALLN-346, reduces hyperuricemia, and uricosuria in urate oxidase-deficient mice. Front Med (Lausanne) 7, 569215 (2020).

20 Szczurek, P. et al. Oral uricase eliminates blood uric acid in the hyperuricemic pig model. PLoS One 12, e0179195 (2017).

21 Terkeltaub, R. Update on gout: new therapeutic strategies and options. Nat Rev Rheumatol 6, 30–38 (2010).

22 Schultz, A. C., Nygaard, P. & Saxild, H. H. Functional analysis of 14 genes that constitute the purine catabolic pathway in Bacillus subtilis and evidence for a novel regulon controlled by the PucR transcription activator. J Bacteriol 183, 3293–3302 (2001).

23 Vargas-Santos, A. B., Peloquin, C. E., Zhang, Y. & Neogi, T. Association of chronic kidney disease with allopurinol use in gout treatment. JAMA Intern Med 178, 1526–1533 (2018).

24 Ichida, K. et al. Decreased extra-renal urate excretion is a common cause of hyperuricemia. Nat Commun 3, 764 (2012).

25 Guo, Y. et al. Inulin supplementation ameliorates hyperuricemia and modulates gut microbiota in Uox-knockout mice. Eur J Nutr 60, 2217–2230 (2021).

26 Zhang, W. et al. Variation of serum uric acid is associated with gut microbiota in patients with diabetes mellitus. Front Cell Infect Microbiol 11, 761757 (2022).

27 Wang, J. et al. The gut microbiota as a target to control hyperuricemia pathogenesis: Potential mechanisms and therapeutic strategies. Crit Rev Food Sci Nutr, 1–11 (2021).

28 Garcia-Arroyo, F. E. et al. Probiotic supplements prevented oxonic acid-induced hyperuricemia and renal damage. PLoS One 13, e0202901 (2018).

29 Yun, Y. et al. Intestinal tract is an important organ for lowering serum uric acid in rats. PLoS One 12, e0190194 (2017).

30 Crook, N. et al. Adaptive strategies of the candidate probiotic E. coli Nissle in the mammalian gut. Cell Host Microbe 25, 499–512 (2019).

31 Behnsen, J. et al. Siderophore-mediated zinc acquisition enhances enterobacterial colonization of the inflamed gut. Nat Commun 12, 7016 (2021).

32 Kurtz, C. B. et al. An engineered E. coli Nissle improves hyperammonemia and survival in mice and shows dose-dependent exposure in healthy humans. Sci Transl Med 11, eaau7975 (2019).

33 Puurunen, M. K. et al. Safety and pharmacodynamics of an engineered E. coli Nissle for the treatment of phenylketonuria: a first-in-human phase 1/2a study. Nat Metab 3, 1125–1132 (2021).

34 Praveschotinunt, P. et al. Engineered E. coli Nissle 1917 for the delivery of matrix-tethered therapeutic domains to the gut. Nat Commun 10, 5580 (2019).

35 Isabella, V. M. et al. Development of a synthetic live bacterial therapeutic for the human metabolic disease phenylketonuria. Nat Biotechnol 36, 857–864 (2018).

36 Danino, T. et al. Programmable probiotics for detection of cancer in urine. Sci Transl Med 7, 289ra284 (2015).

37 Rottinghaus, A. G., Ferreiro, A., Fishbein, S. R. S., Dantas, G. & Moon, T. S. Genetically stable CRISPR-based kill switches for engineered microbes. Nat Commun 13, 672 (2022).

38 Park, Y., Espah Borujeni, A., Gorochowski, T. E., Shin, J. & Voigt, C. A. Precision design of stable genetic circuits carried in highly-insulated E. coli genomic landing pads. Mol Syst Biol 16, e9584 (2020).

39 Huang, Q. et al. A signal sequence suppressor mutant that stabilizes an assembled state of the twin arginine translocase. Proc Natl Acad Sci U S A 114, E1958–E1967 (2017).

40 Bury, S. et al. The probiotic Escherichia coli strain Nissle 1917 combats lambdoid bacteriophages stx and lambda. Front Microbiol 9, 929 (2018).

41 Yim, J. et al. Transcriptional profiling of the probiotic Escherichia coli Nissle 1917 strain under simulated microgravity. Int J Mol Sci 21, 2666 (2020).

42 Soundararajan, M., von Bunau, R. & Oelschlaeger, T. A. K5 capsule and lipopolysaccharide are important in resistance to T4 phage attack in probiotic E. coli strain Nissle 1917. Front Microbiol 10, 2783 (2019).

43 Lu, J. et al. Knockout of the urate oxidase gene provides a stable mouse model of hyperuricemia associated with metabolic disorders. Kidney Int 93, 69–80 (2018).

44 Stamp, L. K. & Dalbeth, N. Prevention and treatment of gout. Nat Rev Rheumatol 15, 68–70 (2019).

45 Maier, L. et al. Extensive impact of non-antibiotic drugs on human gut bacteria. Nature 555, 623–628 (2018).

46 Chu, Y. et al. Metagenomic analysis revealed the potential role of gut microbiome in gout. NPJ Biofilms Microbiomes 7, 66 (2021).

47 Yu, Y., Liu, Q., Li, H., Wen, C. & He, Z. Alterations of the gut microbiome associated with the treatment of hyperuricaemia in male rats. Front Microbiol 9, 2233 (2018).

48 Lin, A. et al. Self-cascade uricase/catalase mimics alleviate acute gout. Nano Lett 22, 508–516 (2022).

49 Ming, J. et al. A novel cascade nanoreactor integrating two-dimensional Pd-Ru nanozyme, uricase and red blood cell membrane for highly efficient hyperuricemia treatment. Small 17, e2103645 (2021).

50 Otani, N. et al. Recent approaches to gout drug discovery: an update. Expert Opin Drug Discov 15, 943–954 (2020).

51 Schlesinger, N., Yasothan, U. & Kirkpatrick, P. Pegloticase. Nat Rev Drug Discov 10, 17–18 (2011).

52 FitzGerald, J. D. et al. 2020 American college of rheumatology guideline for the management of gout. Arthritis Care Res (Hoboken) 72, 744–760 (2020).

53 Litvak, Y. et al. Commensal enterobacteriaceae protect against Salmonella colonization through oxygen competition. Cell Host Microbe 25, 128–139 (2019).

54 Iwadate, Y. & Kato, J. I. Identification of a formate-dependent uric acid degradation pathway in Escherichia coli. J Bacteriol 201, e00573–00518 (2019).

55 Papakostas, K. & Frillingos, S. Substrate selectivity of YgfU, a uric acid transporter from Escherichia coli. J Biol Chem 287, 15684–15695 (2012).

56 Cai, L. et al. Construction and expression of recombinant uricaseexpressing genetically engineered bacteria and its application in rat model of hyperuricemia. Int J Mol Med 45, 1488–1500 (2020).

57 Sarate, P. J. et al. E. coli Nissle 1917 is a safe mucosal delivery vector for a birch-grass pollen chimera to prevent allergic poly-sensitization. Mucosal Immunol 12, 132–144 (2019).

58 Lin, S. et al. Mucosal immunity-mediated modulation of the gut microbiome by oral delivery of probiotics into Peyer’s patches. Sci Adv 7, eabf0677 (2021).

59 Hwang, I. Y. et al. Engineered probiotic Escherichia coli can eliminate and prevent Pseudomonas aeruginosa gut infection in animal models. Nat Commun 8, 15028 (2017).

60 Rao, S. et al. Toward a live microbial microbicide for HIV: commensal bacteria secreting an HIV fusion inhibitor peptide. Proc Natl Acad Sci U S A 102, 11993–11998 (2005).

61 Sassone-Corsi, M. et al. Microcins mediate competition among Enterobacteriaceae in the inflamed gut. Nature 540, 280–283 (2016).

62 Bermudez-Humaran, L. G. & Langella, P. Live bacterial biotherapeutics in the clinic. Nat Biotechnol 36, 816–818 (2018).

63 Ou, B. et al. Engineered recombinant Escherichia coli probiotic strains integrated with F4 and F18 fimbriae cluster genes in the chromosome and their assessment of immunogenic efficacy in vivo. ACS Synth Biol 9, 412–426 (2020).

64 Hu, Y. et al. OmpR positively regulates urease expression to enhance acid survival of Yersinia pseudotuberculosis. Microbiology (Reading) 155, 2522–2531 (2009).

65 Milton, D. L., O’Toole, R., Horstedt, P. & Wolf-Watz, H. Flagellin A is essential for the virulence of Vibrio anguillarum. J Bacteriol 178, 1310–1319 (1996).

66 Datsenko, K. A. & Wanner, B. L. One-step inactivation of chromosomal genes in Escherichia coli K-12 using PCR products. Proc Natl Acad Sci U S A 97, 6640–6645 (2000).

67 Zhu, Y., Peng, X. & Ling, G. An update on the animal models in hyperuricaemia research. Clin Exp Rheumatol 35, 860–864 (2017).

68 de Muinck, E. J., Trosvik, P., Gilfillan, G. D., Hov, J. R. & Sundaram, A. Y. M. A novel ultra high-throughput 16S rRNA gene amplicon sequencing library preparation method for the Illumina HiSeq platform. Microbiome 5, 68 (2017).

