## Supplementary material for "Insulated expression of periplasmic uricase in *E. coli* Nissle 1917 for the treatment of hyperuricemia": Combined supplementary file

[*] For correspondence:

**Supplementary Table**

**Supplementary Table 1.** Strains, plasmids and primers used in this study.

| Name | Description | Source or application |
|---|---|---|
| Strains | | |
| EcN | *E. coli* Nissle 1917 (serotype O6:K5:H1) | Lab collection |
| EcN C6 | Recombinant EcN strain with insulated expression of uricase between *uspG* and *ahpF* genes. | This study |
| S17-1 | *thi pro hsdR*$^{-}$ *hsd*M$^{+}$ *recA* RP4 2-Tc::Mu-Km::Tn*7* | Lab collection |
| DH5α | Strain used for clone construction | Lab collection |
| *Salmonella* Typhimurium LT2 | *Salmonella enterica* serovar Typhimurium LT2 (wild-type *S*. Typhimurium LT2) | Lab collection |
| Plasmids | | |
| pKT100 | Cloning vector, p15A replicon, Kan$^{R}$ | (64) |
| pKT-*ompA-uricase* | Plasmid expressing OmpA-uricase, Kan$^{R}$ | This study |
| pKT-*tamA-uricase* | Plasmid expressing TamA-uricase, Kan$^{R}$ | This study |
| pKT-*ftsP-uricase* | Plasmid expressing FtsP-uricase, Kan$^{R}$ | This study |
| pKT-*lpp-ompA-uricase* | Plasmid expressing Lpp-OmpA-uricase, Kan$^{R}$ | This study |
| pKT-*yebF-uricase* | Plasmid expressing YebF-uricase, Kan$^{R}$ | This study |
| pKT-*inpNC-uricase* | Plasmid expressing InpNC-uricase, Kan$^{R}$ | This study |
| pKT-*ftsP-gfp* | Plasmid expressing FtsP-GFP, Kan$^{R}$ | This study |
| pDM4 | Suicide vector for recombinant strain construction, Cm$^{r}$ | (65) |
| pDM4-*UAm*-C6 | Vector of insertion UAm-C6 fragment | This study |
| Oligonucleotides | | |
| *ftsP*-Forward | CAATTTCACACAAGAAGGAGATCACATATGTCACTCAGTCGGCGTCAG | EcN C6 inserted fragment |
| *ftsP*-Reverse | GTACTTCGTTGTCATTGGATTGCCCGGCTGCGCTGGCCTTC | |
| *uricase*-Forward | TCCAATGACAACGAAGTACCTGGTTCCATGACTGCCACCGCAGAAACCTC | |
| *uricase*- Reverse | GCAGTCGATCGTACGCTACTAGCAGAATCCGGCGATGTTC | |
| *P6*- Forward | AGTATATACACTCCGCTAATGTGAGTTAGCTCACTCATTAGGCACCCCAGGCTTGACA | |
| *P6*- Reverse | CTCCTTCTTGTGTGAAATTGCACACATGCTAGGAGCCGATGATTAATTGTCAAGCCT | |

| | | |
|---|---|---|
| *rrnBT*- Forward | CGAACCTGAACCACTACCATAAAACGAAAGGCCCAGTCTTTCGAC | |
| *rrnBT*- Reverse | TAGCGTACGATCGACTGCCAGGCATCAAATAAAACGAAAGGCTCAGTCGAAAGACTGG | |
| *uspG*- Forward | CTAGCGGAGTGTATATCAAGACCAGCCAGAAGCTGCTGGCGA | pDM4-UAm-C6 construction |
| *uspG*- Reverse | TAGCGGAGTGTATATACTGGCATTAAAAAGCCCTGCAGGGATGGCTCCGG | |
| *ahpF*- Forward | GGTAGTGGTTCAGGTTCGCAATAAAAAAGCCGCCAGGTTTGAC | |
| *ahpF*- Reverse | CAGGTTACCCGCATGCAAGACGACGTTGATGTGATCGACAGC | |
| *gfp*- Forward | TCCAATGACAACGAAGTACCTGGTTCCAGCAAGGGCGAGGAGCTGTTCA | pKT-*ftsP*-*gfp* construction |
| *gfp*- Reverse | CGAACCTGAACCACTACCTTACTTGTACAGCTCGTCCATGCC | |
| *ompA*-Forward | CAATTTCACACAAGAAGGAGATCACATATGAAAAAGACAGCTATCGCGAT | pKT-*ompA*-*uricase* construction |
| *ompA*-Reverse | GTACTTCGTTGTCATTGGAGGTGTTATCTTTCGGAGCGGCCTG | |
| *tamA*-Forward | CAATTTCACACAAGAAGGAGATCACATATGCGCTATATCCAACAGTTAT | pKT-*tamA*-*uricase* construction |
| *tamA*-Reverse | GTACTTCGTTGTCATTGGACTGTAGACGGACGTTCGCGG | |
| *lpp*-*ompA*-Forward | CAATTTCACACAAGAAGGAGATCACATATGAAAGCTACTAAACTGGTACTGG | pKT-*lpp*-*ompA*-*uricase* construction |
| *lpp*-*ompA*-Reverse | CAAAGCCAACATACGGGTTAATTCCCTGATCGATTTTAGCGTTGCTGGA | |
| *yebF*-Forward | CAATTTCACACAAGAAGGAGATCACATATGAAAAAAAGAGGGGCGTTTTTAGGGC | pKT-*yebF*-*uricase* construction |
| *yebF*-Reverse | GTACTTCGTTGTCATTGGAACGCCGCTGATATTCCGCCAT | |
| *inpNC*-Forward | CAATTTCACACAAGAAGGAGATCACATATGACTCTCGACAAGGCGTTGGTG | pKT-*inpNC*-*uricase* construction |
| *inpNC*-Reverse | CCACCACTGCCGCCACCTTTAACCTCGATCCAATCATCATCTTCG | |
| *fimA*-Forward | ATACTACGACGGTAAATGGT | qPCR for examining the copy numbers of EcN C6 |
| *fimA*-Reverse | CGGCTTTTGTGGCAACAGTGG | |

| | | |
|---|---|---|
| | | genome |
| *uox*-Forward 1 | CAACTCAACAGGAAAGAGATCCAG | Identification of C57BL/6J *uox*-knockout mouse |
| *uox*-Reverse 1 | GTGTTGCCGCCATCTCTGCCTCTAGGC | |
| *uox*-Forward 2 | GTAATAACAGGATAGAGTCTCCTCGG | |
| *uox*-Reverse 2 | GGATGAATGCATGGACGTGTTTGATCCC | |

### Supplementary Figures

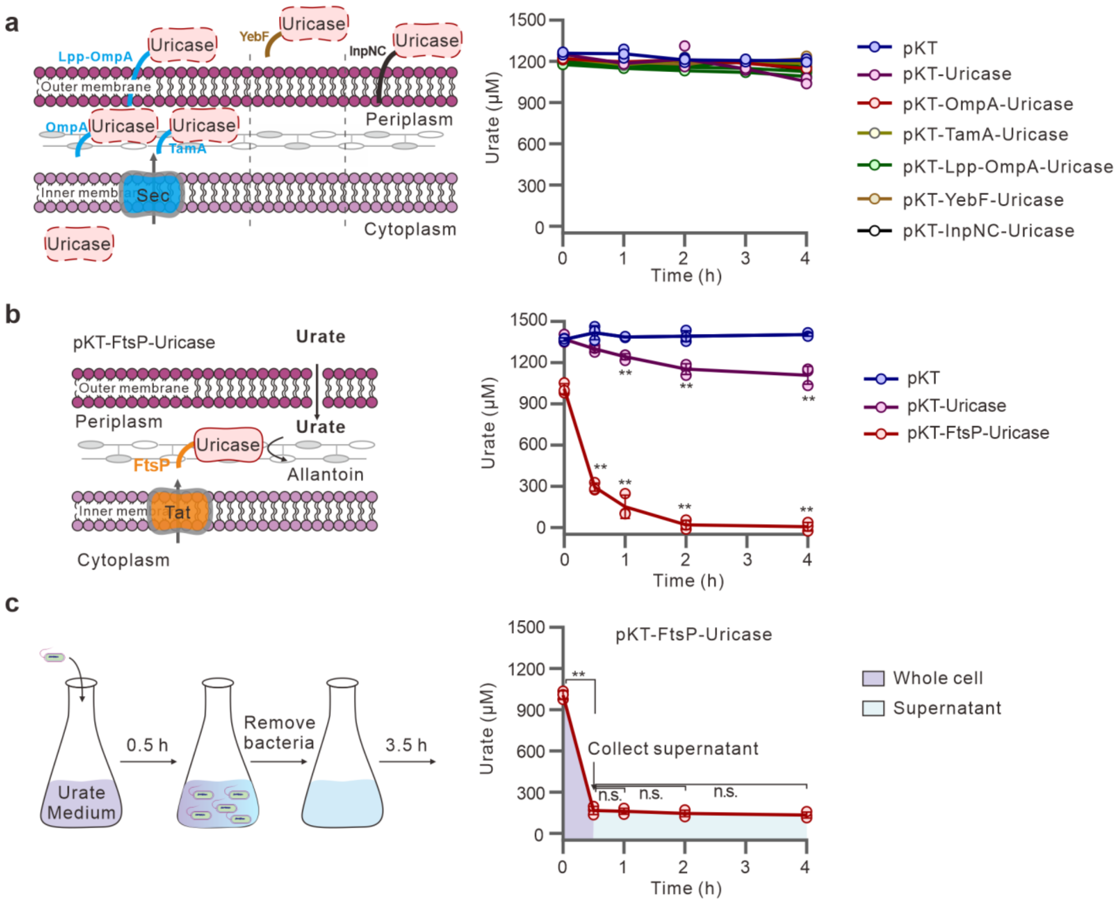


**Supplementary Fig. 1.** Uricase activity of the recombinant strains expressing uricase gene fused with different signal peptides. **a** Overview of the proposed locations of uricase fused with different signal peptides (left) and activities of the corresponding strains in degrading urate *in vitro* (right). **b** Diagram of periplasmic expression of uricase driven by the FtsP peptide through the Tat secretion system (left), and the urate degradation activity of the strains *in vitro* (right). **c** Uricase activity of whole cell or supernatant from the EcN strain expressing FtsP-uricase fused protein. Data were from triplicate independent bacterial cultures; bars indicate the mean ± SD based on two-tailed unpaired Student’s t test. (**P < 0.01).

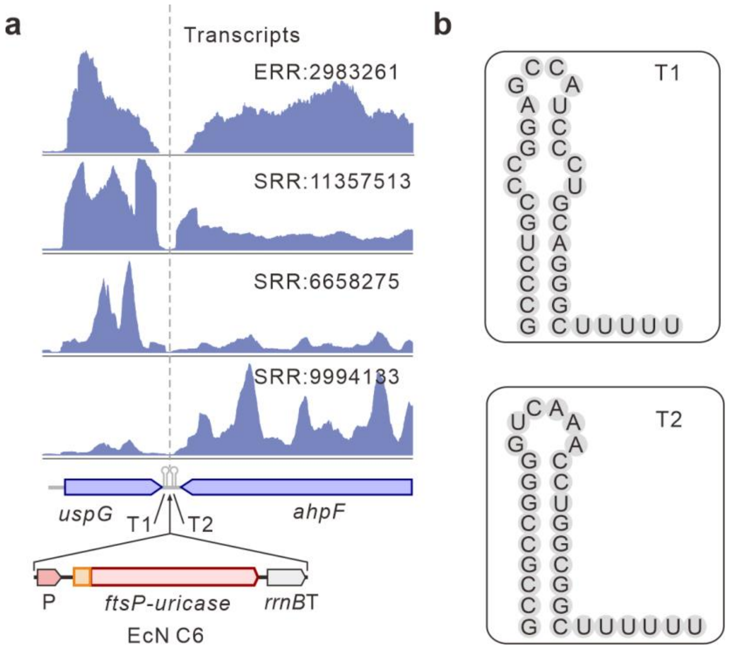


**Supplementary Fig. 2.** Insulated site located between *uspG* and *ahpF* genes in *E. coli*. **a** Normalized RNA levels in regions covering *uspG* and *ahpF* genes from different studies. The site for insertion of the *ftsP-uricase* fragment in the EcN genome is shown at the bottom. **b** RNA secondary structure of the T1 and T2 terminators predicted by RNAfold service (http://beagle.bio.uniroma2.it/).

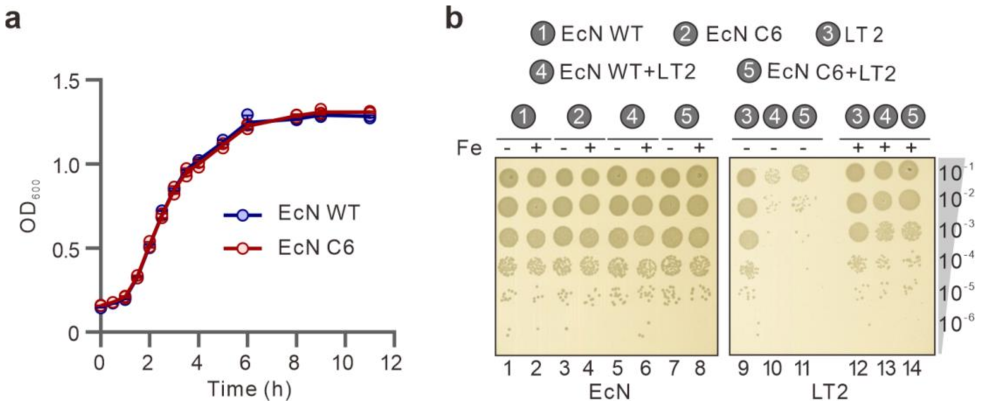


**Supplementary Fig. 3.** Influences of uricase expression on the characteristics of EcN C6. **a** Growth comparison of the EcN WT and the EcN C6 strains. **b** Competitive growth assays of EcN WT, C6 and *S*. Typhimurium LT2.

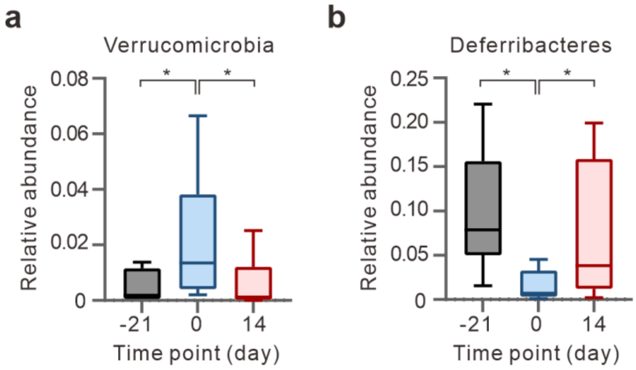


**Supplementary Fig. 4.** Relative abundance of Verrucomicrobia (**a**) and Deferribacteres (**b**) in the gut microbiota of the EcN C6-administrated rats at -21, 0 and 14 days (n = 10). *P < 0.05.

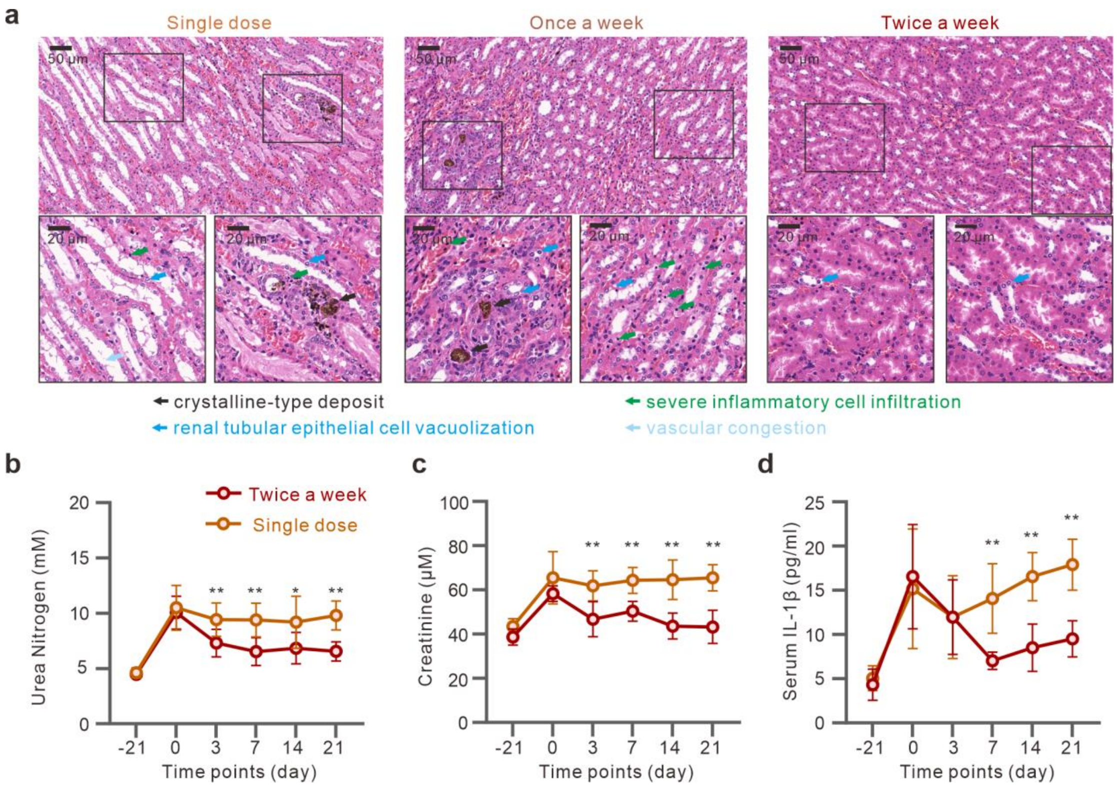


**Supplementary Fig. 5.** Treatment with EcN C6 alleviates kidney damage (**a**) and decreases serum urea nitrogen (**b**), creatinine (**c**), and IL-1β (**d**) levels in hyperuricemia rats. Scale bars: 50 μm or 20 μm; **P < 0.01; *P < 0.05.
